# Identification and characterization of a β-1,4 Galactosidase from *Elizabethkingia meningoseptica* and its application on living cell surface

**DOI:** 10.1101/2023.10.12.561795

**Authors:** Yongliang Tong, Xinrong Lu, Danfeng Shen, Lin Rao, Lin Zou, Shaoxian Lyu, Linlin Hou, Guiqin Sun, Li Chen

## Abstract

The biological function of terminal galactose on glycoprotein is an open field of research. Although progress had being made on enzymes that can remove the terminal galactose on glycoproteins, there is a lack of report on galactosidases that can work directly on living cells. In this study, a unique beta 1,4 galactosidase was isolated from *Elizabethkingia meningoseptica* (*Em*). It exhibited favorable stability at various temperatures (4-37℃) and pH (5-8) levels and can remove β-1, 4 linked galactoses directly from glycoproteins. Using Alanine scanning, we found that two acidic residues (Glu-468, and Glu-531) in the predicted active pocket are critical for galactosidase activity. In addition, we also demonstrated that it could cleave galactose residues present on living cell surface. As the enzyme has a potential application for living cell glycan editing, we named it glycan editing galactosidase I or geGalaseI. In summary, our findings lay the groundwork for prospective investigations by presenting a prompt and gentle approach for the removal of galactose moieties from cell surface.

## Introduction

β-Galactosidase (EC 3.2.1.23) is an important biocatalyst with both transglycosylation and hydrolysis activities[1]. The first galactosidase was reported in 1889[2], and the first sequence and crystal structure of galactosidase was published in 1983 and 1994 respectively[3, 4]. Galactosidase is widely presented in bacteria, yeast, fungi, plant and mammals[5]. Based on the similarity of protein sequences, β-galactosidases can be classified into 4 glycoside hydrolase (GH) families, including GH-1, GH-2, GH-35, and GH-42[6]. The substrates for galactosidase is the non-reducing β-D-galactose on oligosaccharides and glycans, such as lactose, galacto-oligosaccharides (GOS), free glycans and glycans on glycoproteins[7]. Galactosidase has been widely used in the food industry, especially for the dairy products. Galactosidase treated milk is a solution for lactose intolerant people[8–11]. The galactosidase-generated GOS is an important prebiotic ingredient in dairy products[12–15].

Although N-glycosylation is a common post-translational modification (PTM) observed in all eukaryotic cells[16], report on galactosidase directly on glycoproteins is rare. Butler *et al.* firstly applied bovine testes β-galactosidase in detailed structure analysis for free N-glycans in 2003[17]. Subsequently the method was used for diagnosis of congenital disorders of glycosylation (CDG) and identification of antibody glycoforms[18, 19]. A limitation of the reported method is that N-glycans must be released by PNGaseF from denatured proteins to be a substrate for galactosidase. In 2018, Butler *et al.* reported that a galactosidase from *Bacteroides fragilis* could remove terminal galactose directly from naïve antibody thus could be applied for high yields of single glycoform antibodies[20, 21]. However, all the previous studies on galactosidase activity were conducted in a non-cell system. It is noteworthy that a significant proportion of cell surface proteins are glycoproteins[22, 23]. Some N-glycans on cell surface glycoproteins contain terminal galactose[24].

Studies on galectins have illustrated that galactose on cell surface may play a critical biological function[25, 26]. Galectins could bind to the galactose molecules present on cell surface to regulate cell viability, function, proliferation, and differentiation[27–34]. Nevertheless, research on the function of cell surface galactose is limited partially because of the lack of tool to cleave galactose on living cells.

In this investigation, we isolated a galactosidase from *Elizabethkingia meningoseptica*, demonstrated its activity directly on terminal galactose on living cell surface, conducted preliminary study on its regulatory action on targeted cells, and named it glycan-editing galactosidase I or geGalaseI.

## Experimental procedures

### Bioinformatics analysis of geGalaseI - the putative β-1,4 galactosidase from *E. meningoseptica*

The gene of geGalaseI was obtained from the genomic DNA of FMS-007, one clinically isolated *E. meningoseptica* strain[35]. Other galactosidases in the multiple-sequence alignment were searched in the Carbohydrate-Active enZYmes Database (CAZy), and their protein sequences were downloaded from GenBank™. The identity and similarity of different galactosidases’ sequence were aligned using the sequence align tool from the UniProt website (www.uniprot.org/align), and the final alignment result was exported by Easy Sequencing in PostScript (ESPript3.0) Web site. The structure prediction of geGalaseI was done by the alphafold2 model (based on Google ColabFold). The molecular docking and results presentation is performed using the PyMOL software.

### Cloning and protein expression

The target gene was amplified by polymerase chain reaction (PCR). Recombination reaction was performed by mixing the linearized pET28a vectors and PCR products and incubating with Exnase II from ClonExpress II One Step Cloning kit (Vazyme, #C112). All geGalaseI mutants were generated by the reverse complement primers for PCR with the template of pET28a-geGalaseI (WT) plasmid. The Galectin1 and Endo F3 gene were synthesized and cloned to pET28a plasmid by GenScript Corporation. The constructed pET28a plasmid was transformed into *E. coli* BL21 (DE3) (Yeasen, #11804) separately for protein expression. Finally, all proteins were purified using Ni Sepharose™ 6 Fast Flow (Cytiva, #175318). Endotoxins were removed using a high-capacity endotoxin removal spin kit (Thermo Fisher Scientific, #88275)

### Enzymatic assay against p-nitrophenyl glycosides

For specificity assay, various p-nitrophenyl α/β-glycosides purchased from Carbosynth were selected as substrates. The reaction solution was incubated at 37℃ for 1 h, and then terminated by Na_2_CO_3_. The absorbance at 405 nm (A_405_) was determined. The optimum pH and pH stability of enzyme was investigated in the pH range of 3.0-9.0 using Glycine-HCl buffer (50 mM, pH 3.0-5.0), MES buffer (100 mM, pH 6.0), HEPES buffer (100 mM, pH 7.0) and Tris-HCl buffer (50 mM, pH 8.0-10.0). For optimum pH test, substrate and geGalaseI were mixed with buffers of different pH. The mixtures were incubated at 37℃ for 5 min, and then terminated by addition of 1 M Na_2_CO_3_. For pH stability test, geGalaseI was pre-incubated in different pH buffers without substrate for 30 min at 37℃. And then the pre-incubated solutions were transported into pH-optimum buffer in the presence of substrate and the mixtures were incubated at 37℃ for 5 min. After terminating the reactions, A_405_ was determined. The optimum temperature of enzyme was examined by setting the reaction systems consisted of substrate, geGalaseI and pH-optimum buffer at 4, 20, 30, 37, 45, 55, 65 and 75℃, separately. After incubation for 5 min, the reactions were terminated. For the detection of thermostability data, geGalaseI was pre-treated under different temperatures, and then pH-optimum buffer and substrate were added to incubate for 5 min at optimum temperature. The values at A_405_ were read after termination the reaction with 1 M Na_2_CO_3_. To test the influence of metal ions and compounds on enzyme activity, a series of metal ions (Cu^2+^, Ca^2+^, Fe^2+^, Mg^2+^, Zn^2+^, Ni^2+^ and Mn^2+^) were added to the reaction systems. The values of kinetic constants were calculated via the Michaelis-Menten equation. All reactions were triplicated for statistical evaluations.

### Enzyme activity against lactose

For mass-spectrometry analysis, a 10 μL reaction mixture containing enzyme, lactose and buffer, was prepared and incubated at 37°C for 2 h. Then 200 μL of deionized water was added for dilution. The diluted mixture was injected into a TSQ Quantiva^TM^ triple quadrupole mass spectrometer system (Thermo Fisher Scientific) for ESI-MS (electrosprayionization mass spectrometer) analysis. The reaction product was scanned from m/z 100 to 600 for 1 min in a positive mode.

### Enzyme activity against glycoproteins

To verify the activity of the candidate enzyme, IgG with complex N-glycan, was selected as the standard substrate glycoprotein and subjected to cleavage using a ‘Cut-Heat-Cut’ strategy[36]. In brief, a totally 200 μL reaction system consisted of 100 μg IgG, enzyme reaction buffer and geGalaseI was prepared. Then the mixture was incubated at 37℃ for 12 h and then denatured at 98℃ for 10 min. 5 μL PNGase F was added for releasing all N-linked oligosaccharides from IgG. Finally, the glycans were analyzed by Matrix-Assisted Laser Desorption/Ionization Time-of-Flight (MALDI-TOF).

### Enzyme activity on immune cell surface

To study the enzymatic activity of geGalaseI on living cell, we separated splenic cell from adult C57BL/6 mice. All animal procedures were approved by the Ethics Committee of Experimental Research, Fudan University Shanghai Medical College. Briefly, mice were sacrificed, spleens were extracted and ground on a 200-mesh strainer to obtain a single-cell suspension. After centrifugation, the precipitate was resuspended with erythrocyte lysate (Yeasen, #40401) and left at room temperature for 10 minutes to remove the erythrocytes. The remaining cells were counted and diluted with RPMI-1640 medium to a density of 5 × 10^6^ cells/mL. The obtained cells were subjected to treatment with varying doses of glycosidase for a specific duration.

After centrifugation to remove the enzymes, the galactose on the cell surface was labeled with Galectin 1, detected with Alexa Fluor®488 Conjugated anti-His Antibody (Cell Signaling Technology, #14930), and finally the cell was detected by flow cytometry (BD, FACSCalibur). To observe the glycomics change on cell surface, we treated the cell with Endo F3 (100 μg/mL) to release N-glycans. After 1h incubation, cells were removed by centrifugation. The supernatant was collected and ultrafiltered with a 3 kD ultrafiltration tube to remove the Endo F3. The flow-through liquid was collected for glycan enrichment using a solid phase extraction column (ENVI-Carb). After vacuum dried, the sample was detected by MALDI-TOF.

### Splenic cells and B16-OVA cells coculture

Splenic cells, obtained from OT-I transgenic mice (courtesy of Likun Gong from Shanghai Institute of Material Medica, Chinese Academy of Science), were subjected to a 3-hour treatment with SiaT and geGalaseI. The determination of cell viability was conducted using the 7-AAD kit (Yeasen, #40745). The cells that underwent treatment were subjected to centrifugation in order to exclude enzymes and subsequently resuspended in 1640 medium at a density of 3 × 10^6^ cells/mL. The aforementioned cells were subjected to co-cultivation with B16-OVA cells (at a concentration of 3×10^5^ cells/mL) that had been seeded onto a 96-well plate the day prior. After a 24-hour incubation period, the immune cells that were suspended in the culture medium were carefully extracted for the purpose of conducting an apoptosis assay (Yeasen, #40311).

## Results

### Bioinformatics analysis of a candidate Galactosidase gene in the whole-genome sequence of E. meningoseptica

A clinical strain of *E. meningoseptica*, FMS-007, was isolated from a T-cell non-Hodgkin’s lymphoma patient by our lab. Bioinformatics analysis of the whole genome led to the identification of 5 candidate glycoprotein galactosidases, which were numbered *Em*2006, *Em* 2131, *Em* 2923, *Em* 3018, and *Em* 3052, respectively. After expression purification and activity screening, *Em* 3052 was selected as the subsequent research protein and named with geGalaseI. Phylogenetic analysis of geGalaseI with other galactosidases published in GenBank™ revealed that geGalaseI belonged to the GH2 class of Glycoside Hydrolases (GHs) defined in the CAZy database (Fig. 1). Those characterized galactosidases in GH2 were all β-galactosidase. geGalaseI could be divided into 3 parts, the N-terminal domain, the middle domain and the C-terminal domain (Fig. S1).

**Figure 1.**
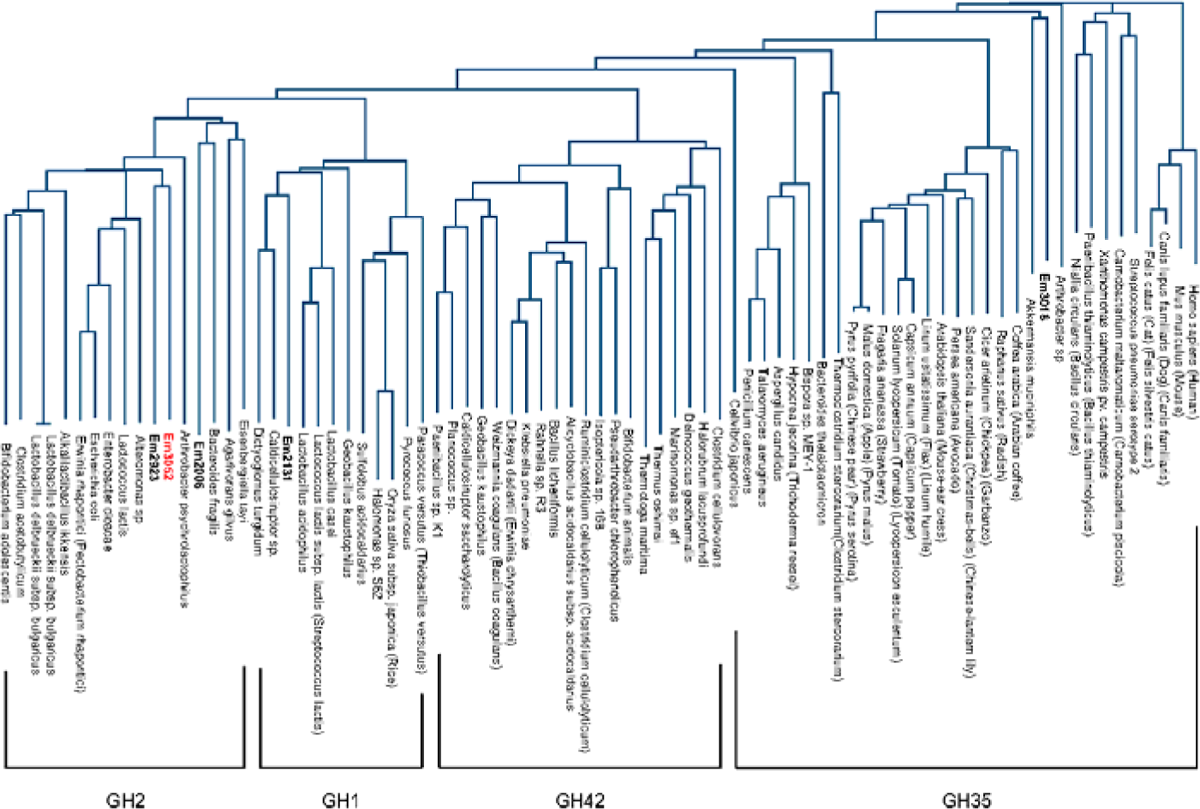
Phylogenetic tree of Candidate galactosidases from *Elizabethkingia meningoseptica* and characterized. β**-galactosidase.** The amino acid sequences of all enzymes were obtained from uniprot Website. Sequence alignment was performed using UniProt align tool. The tree was constructed using a neighbor-joining method. Candidate gene are shown in bold, which are *Em*2006, *Em* 2131, *Em* 2923, *Em* 3018, *Em* 3052. *Em* 3052 was subsequently named geGalaseI and is shown in red.

### Substrate specificity and galactosidase activity of geGalaseI on pNPs

For substrate specificity assay, the enzymatic activity of geGalaseI against 16 p-nitrophenyl α/β-glycosides was tested. Only pNP-β-D-Gal could be hydrolyzed by geGalaseI, indicating that geGalaseI was an β galactosyl-specific glycoside hydrolase (Fig. 2a). The Michaelis constant K_m_ and the maximal velocity V_max_ values were determined from the Michaelis-Menten equation (Fig. 2b). The K_m_ was 0.9010 mM and the V_max_ was 4.781 mM/s. It was measured by dilution that 1.27 μg of geGalaseI can catalyze the conversion of half of substrate (0.03 μmol) in 1 min (Fig. 2c). The specific enzyme activity of geGalaseI was calculated to be 23.6 U/mg.

**Figure 2.**
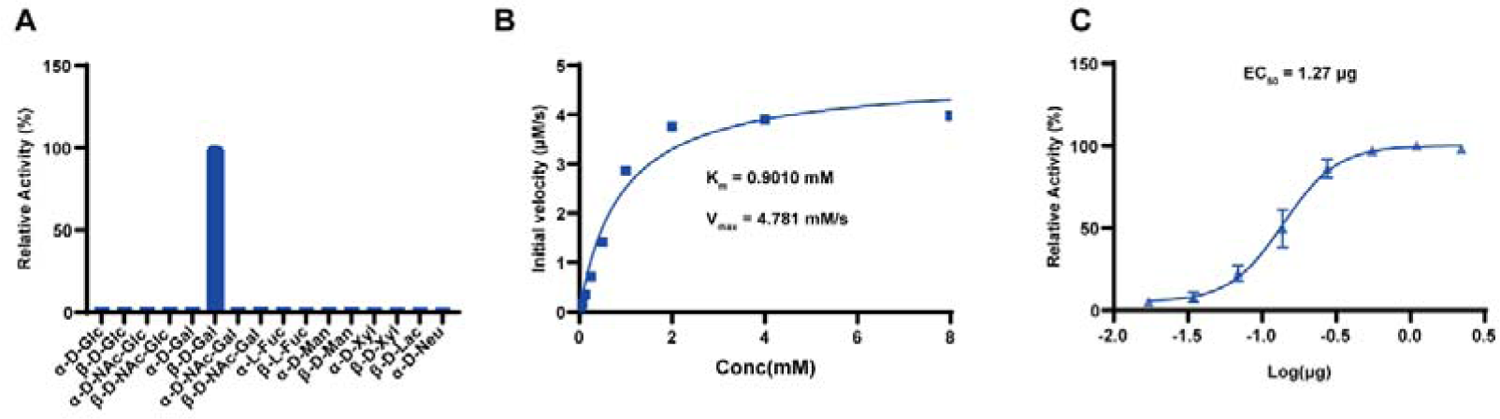
Characterization of geGalaseI. (A) The specificity was shown by testing enzymatic activity of geGalaseI against 16 p-nitrophenyl α/β-glycosides. (B) Michaelis-Menten curve of geGalaseI was shown and K_m_ and V_max_ were calculated to be 0.9010 mM and 4.781 mM/s respectively. (C) The specific enzyme activity of geGalaseI was determined by dilution of geGalaseI and calculated to be 23.6 U/mg.

The optimal temperature for geGalaseI activity was 37℃ (Fig. 3a). Thermostability assay showed when the temperature was between 4 and 37℃, the enzyme activity of geGalaseI was stable. When the temperature was 45℃, the enzyme activity sharply decreased, and when the temperature was 55℃, the enzyme activity was completely lost (Fig. 3b). The optimal pH for geGalaseI activity was between 5 and 8. When the pH was below 4, the activity of geGalaseI was nearly completely lost (Fig. 3c). The pH stability assay showed that at pH values higher than 8, the activity of geGalaseI decreased rapidly. And in the buffer at pH 10, geGalaseI was completely inactivated (Fig. 3d).

**Figure 3.**
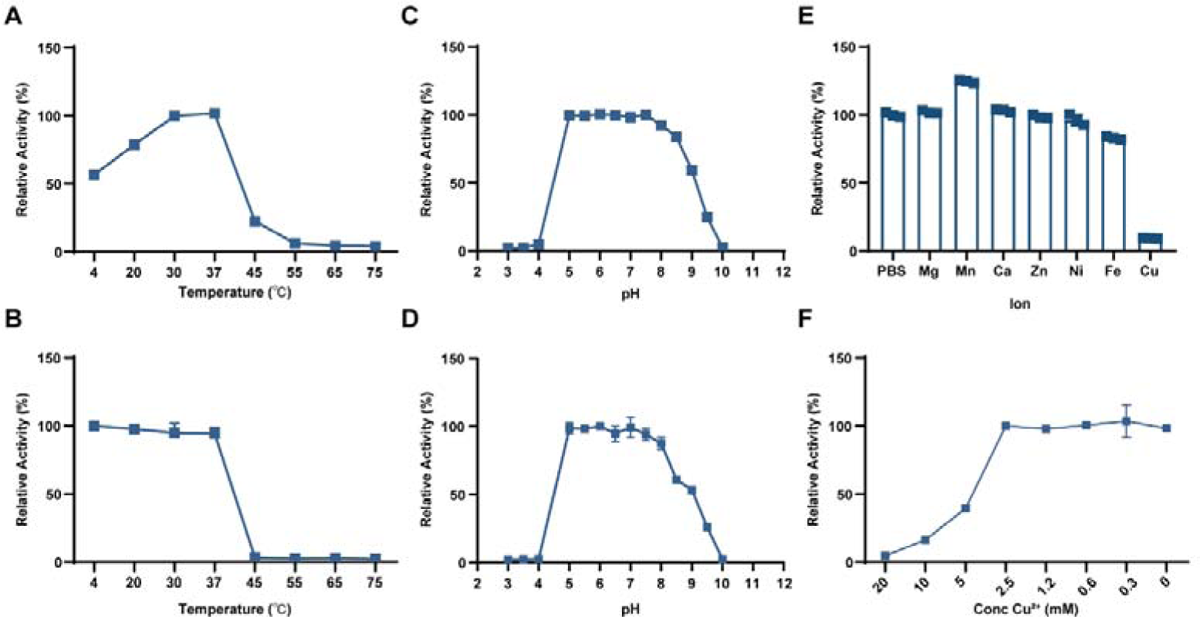
Stability of geGalaseI. (A) The optimal temperature of geGalaseI. (B) The tolerant temperature of geGalaseI. (C) The optimal pH of geGalaseI. (D) The tolerant pH of geGalaseI. (E) Effect of variety of ions on geGalaseI activity. (F) Effect of concentration of Cu^2+^ on geGalaseI activity.

The test of the effect of metal ions on geGalaseI showed that at the final concentration of 20 mM, Cu^2+^ had obvious inhibition effect on the enzyme activity while Mg^2+^, Fe^2+^, Zn^2+^, Ni^2+^ and Mn^2+^, Ca^2+^ had no effects on the enzyme activity of geGalaseI. (Fig. 3e). And the inhibitory effect on geGalaseI activity disappeared when the concentration of Cu^2+^ was less than 2.5 mM (Fig. 3f).

### Defining the enzymatic active sites of geGalaseI

The sequences of geGalaseI and 7 characterized galactosidases (mainly β galactosidase activity) from the GH2 family in the CAZy database were aligned to find relatively conserved sites (Fig. S2). It could be seen from the multiple sequence alignment result that Asn107, Asp211, Asn600, and Trp1005 (marked as green solid pentagram below sequences) of geGalaseI were highly conserved or completely conserved, corresponding to the binding sites of LacZ from *Escherichia coli strain K12*. It suggested that these 4 amino acid sites might be the potential binding sites of geGalaseI. Also, in LacZ, Glu462 and Glu538 acted as catalytic acid and catalytic base, respectively. It could be seen that these two catalytic sites were completely conserved, corresponding to Glu468 and Glu531 of geGalaseI, respectively (marked as green solid triangle below sequences). In addition, His363 and His397 can act as stabilizers of the transition state, which are also conserved in geGalaseI (marked as green hollow pentagram below sequences).

The result of structure prediction of geGalaseI using PyMOL software showed that binding sites and catalytic sites were so close that they might form an active pocket (Fig. 4a). Except Trp1005, other binding sites and catalytic sites were all located in the middle domain of geGalaseI. To verify the predicted active sites by the sequence alignments, a series of mutants of geGalaseI were prepared. The enzymatic activity of the mutants was verified using pNP-β-Gal as a substrate. The results showed that mutant activity of D211, H397, E468, E531, N600, W1005 activity was largely or completely lost, while the mutant activity of N107 and H363 was not impaired (Fig. 4b).

**Figure 4.**
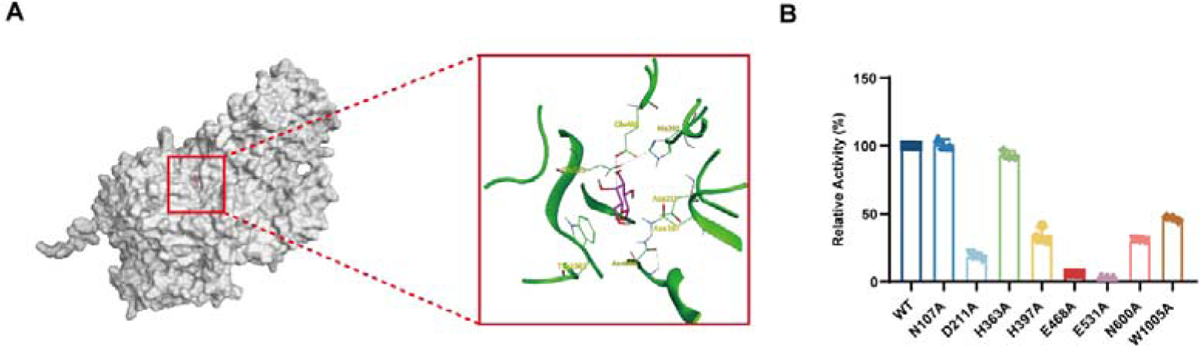
The enzymatic active sites of geGalaseI. (A) Molecular docking of predicted structure of geGalaseI and β galactose was performed using PyMOL software. The left picture is the overall prediction result and the right side is the local zoomed-in image. (B) Enzyme activity aganst pNP-β-D-Gal of 8 mutants of geGalaseI were tested.

### Enzymatic characterization of geGalaseI with lactose

The glycosidic bond specificity of geGalaseI was examined by treating lactose. ESI-MS results showed that in the control group, the ion peak at m/z 365.11 represented a sodium adduct of Gal1-β-4-Glc. In the geGalaseI-treated group, the ion peak at m/z203.05 represented the peak of sodium adduct of single Galactose or Glucose (Fig. 5a). The above mass spectrometry results indicated that geGalaseI could hydrolyze the β1,4 glycosidic bond in lactose.

**Figure 5.**
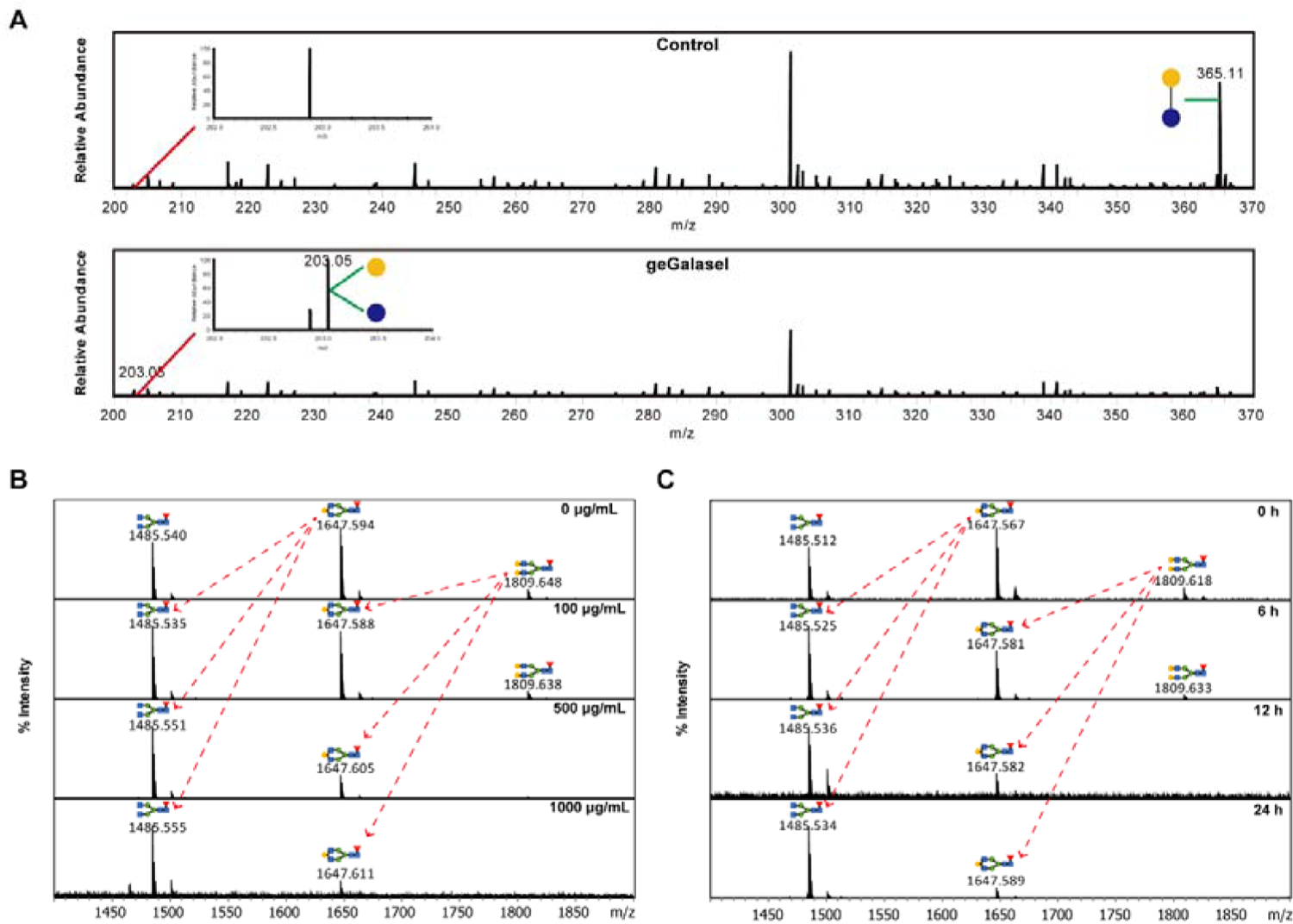
Enzymatic characterization of geGalaseI with oligosaccharides and IgG. (A) Lactose was incubated in the absence (upper) and presence (lower) of geGalaseI, and the reaction mixtures were subjected to ESI-MS analysis. The molecular ion peaks at m/z 365.11 correspond to sodium adducts of Lactose. The molecular ion peaks at m/z 203.05 was sodium adducts of galactose or glucose. (B & C) Serial concentrations of geGalaseI were co-incubated with IgG for different times. The N-glycan were released by PNGaseF and detected by MALDI-TOF. All the peaks were annotated with GlycoMod server and referenced to the previous study; all molecular ions are present in sodiated form ([M + Na]^+^).

### Enzymatic assay of geGalaseI against glycoprotein IgG

IgG produced by CHO cell line was mainly modified by 3 types of complex N-glycan including G0F, G1F, and G2F[37]. To verify the hydrolysis activity of geGalaseI on the β-1,4 galactose of IgG, MALDI-TOF analysis was performed. Results in MALDI-TOF analysis showed that after the treatment of geGalaseI, ion peaks of the sodium adduct of G2F (m/z 1809.648) disappeared completely and that of G1F (m/z 1647.594) decreased dramatically. The decrease of G1F was positively correlated with the concentration and treatment time of geGalaseI. And the main glycan ion peak in geGalaseI-treated group was a sodium adduct of G0 (m/z 1485.553) (Fig. 5b & 5c). These results illustrated that geGalaseI could remove terminal galactose from native antibodies.

### Capacity of geGalaseI to cleave terminal galactose on cell surface

Prior research has indicated that the predominant kind of glycans found on the surface of immune cells are complex glycans[23, 24]. The cleavage of geGalaseI to β-1,4 galactose may be prevented in the presence of α-linked sialic acid (Fig. 6a). In order to confirm the hydrolysis activity of geGalaseI against the terminal galactose on cell surface, a sialidase named SiaT was employed to eliminate α-linked sialic acid (data not shown). Since sialic acid is negatively charged[38], there lacks glycan ion peak of glycoform with sialic acid in the mass spectrum in the positron mode (data not shown). As shown in Fig 6b, the G2 glycoform of exposed galactose was showed on the mass spectrometry graph after SiaT removed the terminal sialic acid. When treated with combination of SiaT and geGalaseI, peaks G1 and G0 glycoforms appeared, indicating that geGalaseI can cleave galactose on the cell surface (Fig. 6b).

**Figure 6.**
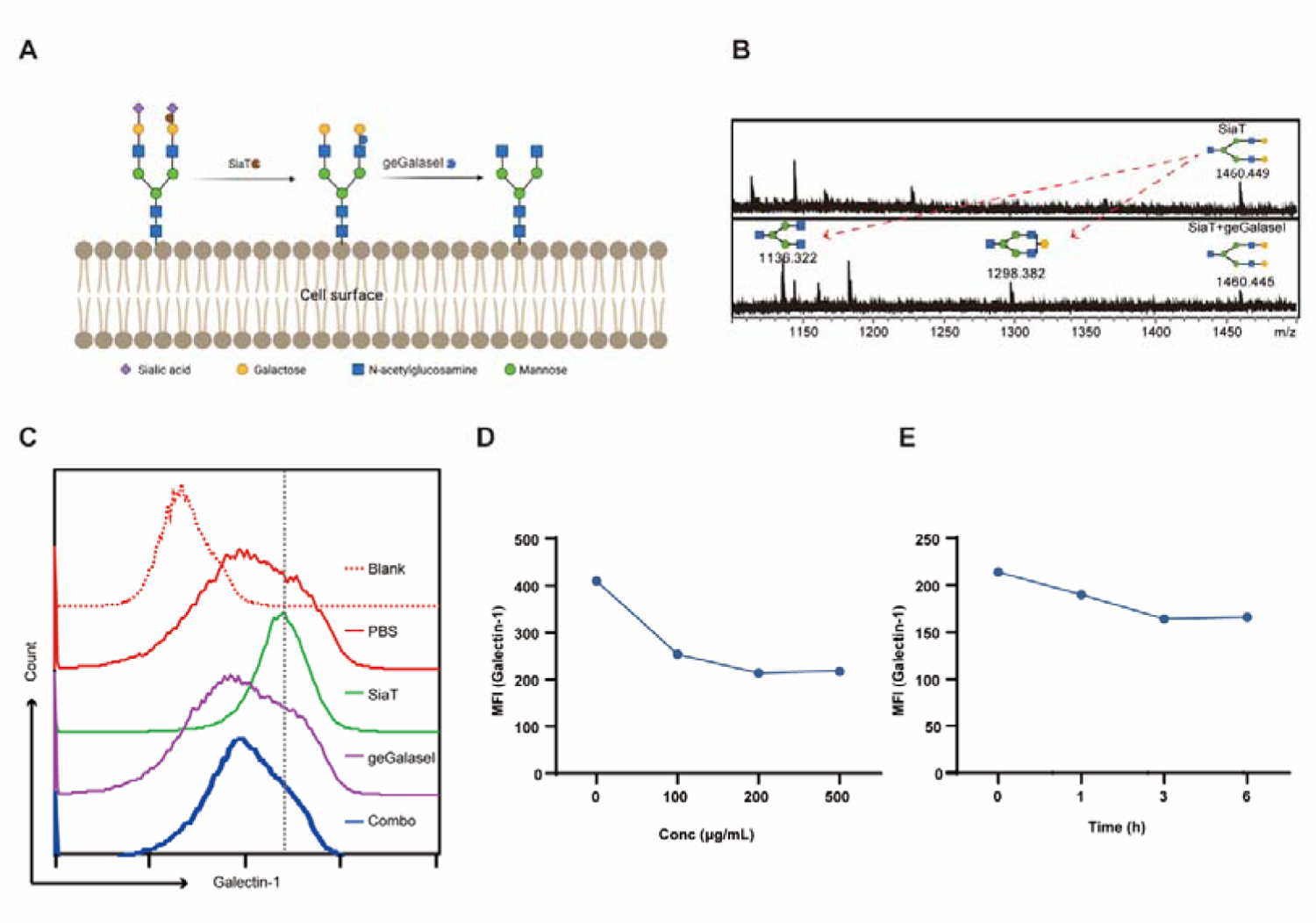
Enzymatic activity of geGalaseI on the surface of living cells. (A) Schematic representation of the changes in cell surface glycans after treatment with SiaT and geGalaseI. (B) Complex N-glycans on the cell surface were released by EndoF3 (100 μg/mL) and then the enriched N-glycans were purified and detected by MALDI-TOF. The top of the graphs is SiaT-treated and the bottom is co-treated. All the peaks were annotated with GlycoMod server and referenced to the previous study. (C) The graph shows the effect of different treatments on the fluorescence intensity of Galectin1 binding on the cell surface as determined by flow cytometry. (D & E) Changes in median fluorescence intensity (MFI) of Galectin1 with geGalaseI concentration and treatment time were shown.

Galectin1 are reported to recognize the carbohydrate motif with lactose or poly-N-acetyllactosamine[27]. Flow cytometry results showed increased MFI (mean fluorescence intensity) of Galectin1 after SiaT treatment, indicating that more galactose residues were exposed. When combined geGalaseI with SiaT, the MFI of Galectin1 was observed to revert back to pre-SiaT level. This observation suggests that geGalaseI effectively removed the galactose residues that were exposed by SiaT treatment, as depicted in Fig. 6c. The optimal timing (3h) and concentration (200 μg/mL) of geGalaseI treatment were determined through Galectin 1 labeling, as shown in Fig. 6d and Fig. 6e.

### Impact of geGalaseI treatment on splenic cells

In the aforementioned experiment, we observed a decrease in Galectin1 binding to cells after galactose removal (Fig. 6c). Galectin1 was reported to be secreted by tumor cell and could cause apoptosis of immune cells[39]. To investigate the impact of alterations in the glycan composition of the cellular membrane on immune response, we conducted a co-culture experiment involving B16-OVA cells and splenic cells obtained from OT-I mice. Cell viability was firstly assessed following SiaT and geGalaseI treatment, and it was seen that the treatment did not result in any impairment of immune cells (Fig. 7a). Next, we observed the apoptosis of splenocytes after co-culture. Interestingly, SiaT treatment led to an increase in apoptosis of splenic cells whereas geGalaseI treatment could rescue the apoptosis caused by SiaT (Fig. 7b). These results are consistent with our observation of Galectin1 binding.

**Figure 7.**
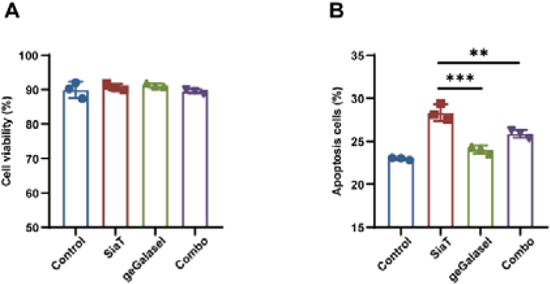
Impact of geGalaseI treatment on splenic cell viability and function. (A) Splenocytes were treated with 200 μg/ml of enzyme for 3h and then stained with 7-AAD and examined by flow cytometry. (B) Splenic cell from OT-I mice was pre-treated with enzyme and then incubated with B16-OVA cell. After 24 h coculture, the splenic cells were collected and stained with Annexin V and PI for apoptosis detection. Data are presented as mean ± standard deviation (SD); **p< 0.01, and ***p< 0.005 using one-way analysis of variance (ANOVA) with Tukey’s multiple comparison test.

## Discussion

β-galactosidase has been discovered for more than a century and its biochemical rection and biological function have been studied in detail[40]. Nevertheless, researches on galactosidase activity still stayed at a cell free level and studies on its impact on cell regulation is limited[41]. In the present study, we isolated a novel β-galactosidase that can act directly on substrates on cell surface from *Elizabethkingia meningoseptica* and named it as glycan editing galactosidase I (geGalaseI). The stability and activity of the geGalaseI were assessed, and the active site of geGalaseI was identified through point mutation. Most importantly, it was determined that geGalaseI possesses the capability to directly cleave galactose moieties present on the surface of living cells. Additionally, we observed that the removal of galactose resulted in a decrease of Galactin1 binding and a reduction of apoptosis of splenic cells. Our findings are consistent with Galectin1 blocking assay, and the molecular mechanism by which removal of galactose leads to reduced apoptosis requires more research[42, 43]. Glycans on cell surface, including the ones with terminal galactose, may have new and/or uncharacterized biological function with significant impacts on basic and translational studies[16]. These cell surface glycans provide an additional molecular layer that facilitates complex cellular interactions with environmental factors[24].

Hence, the structural regulation of cell surface glycans, or cell surface glycan editing, provides a new way for our understanding of glycoscience and translational glycobiology[44]. Current glycan editing is mainly achieved by genetic manipulation and chemoenzymatic treatment [45, 46]. Although genetic manipulation of key enzymes involved in glycan processing is a practical way of glycan editing, its application is rare for long-term side effects [47, 48]. Bertozzi *et al*. used sialidase to excise cell surface sialic acid and studied its biological function[49, 50]. Huang *et al.* employed a pair of Endo F3 with hydrolysis and synthetic activities respectively to replace all N-glycans on cell surface with an N-glycan synthesized in vitro[51]. In addition to sialic editing and global replacement, this study reported a terminal galactose editing for native N-glycans on cell surface.

Cell therapy such as CAR (Chimeric Antigen Receptor)-T cell therapy and stem cell therapy has emerged as a novel approach for unmatched clinical needs[52, 53]. The maintenance and improvement of cell activity in vivo is an important challenge for cell therapy[54]. Previous studies have demonstrated that cellular viability may significantly influenced by the interaction between galactose present on cell surface and galectins[26, 28, 30]. Apparently, altering the galactose composition on cell surface is an alternative way for galectin-based applications. The geGalaseI reported in this study may present a practical way for this approach.

## Notes

### Competing Interest Statement

The authors have declared no competing interest.

