## Supplementary material for "Identification and characterization of a β-1,4 Galactosidase from *Elizabethkingia meningoseptica* and its application on living cell surface": Figures

**Supplementary**
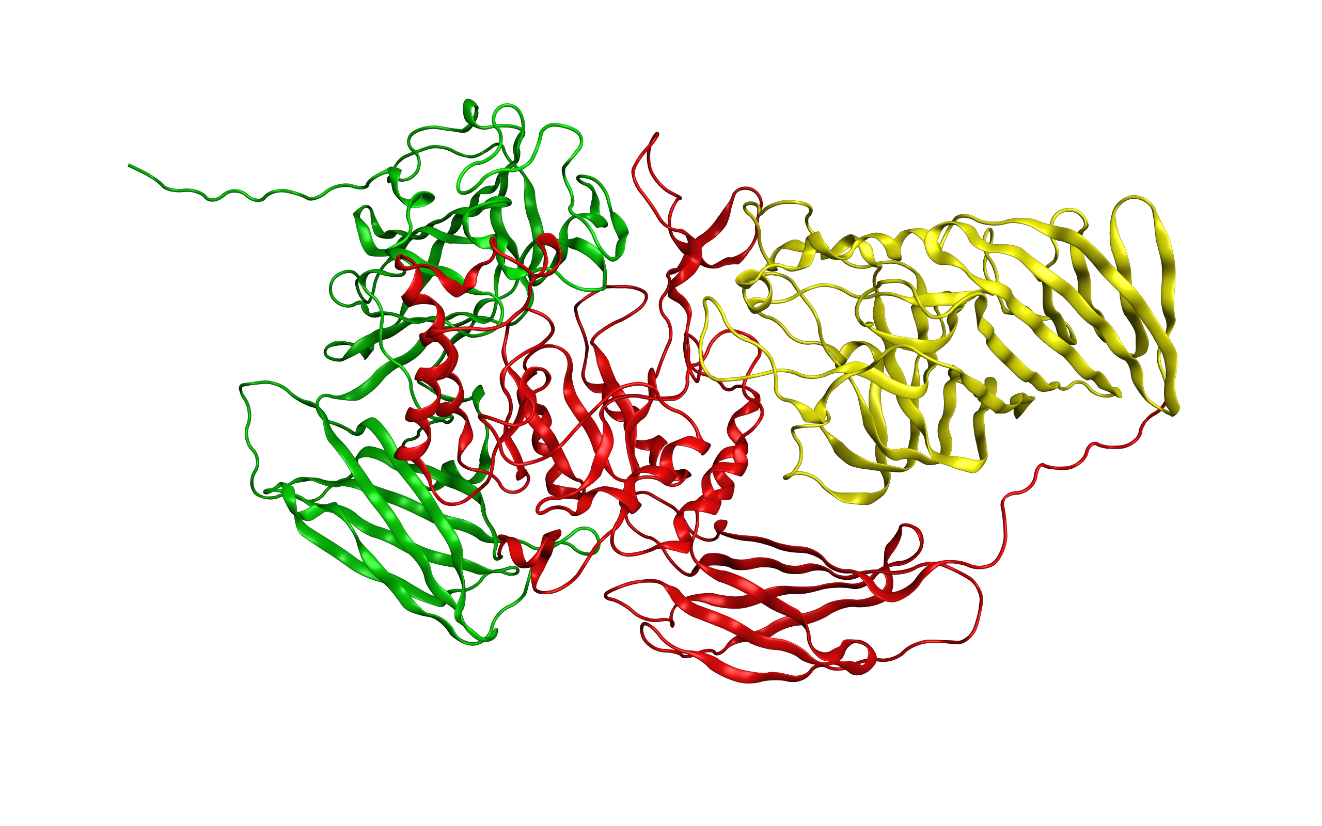

**Fig. S1 Predicted ribbon structure of geGalaseI by Alphafold2.** The 1030 amino acid of geGalaseI was input in to alphafold2 model (based on Google ColabFold) and the structure was displayed by PyMOL. The N-terminal domain (1-304 aa) was colored in green, middle domain (305-737 aa) was in red and the C-terminal domain (738-1030 aa) was in yellow.

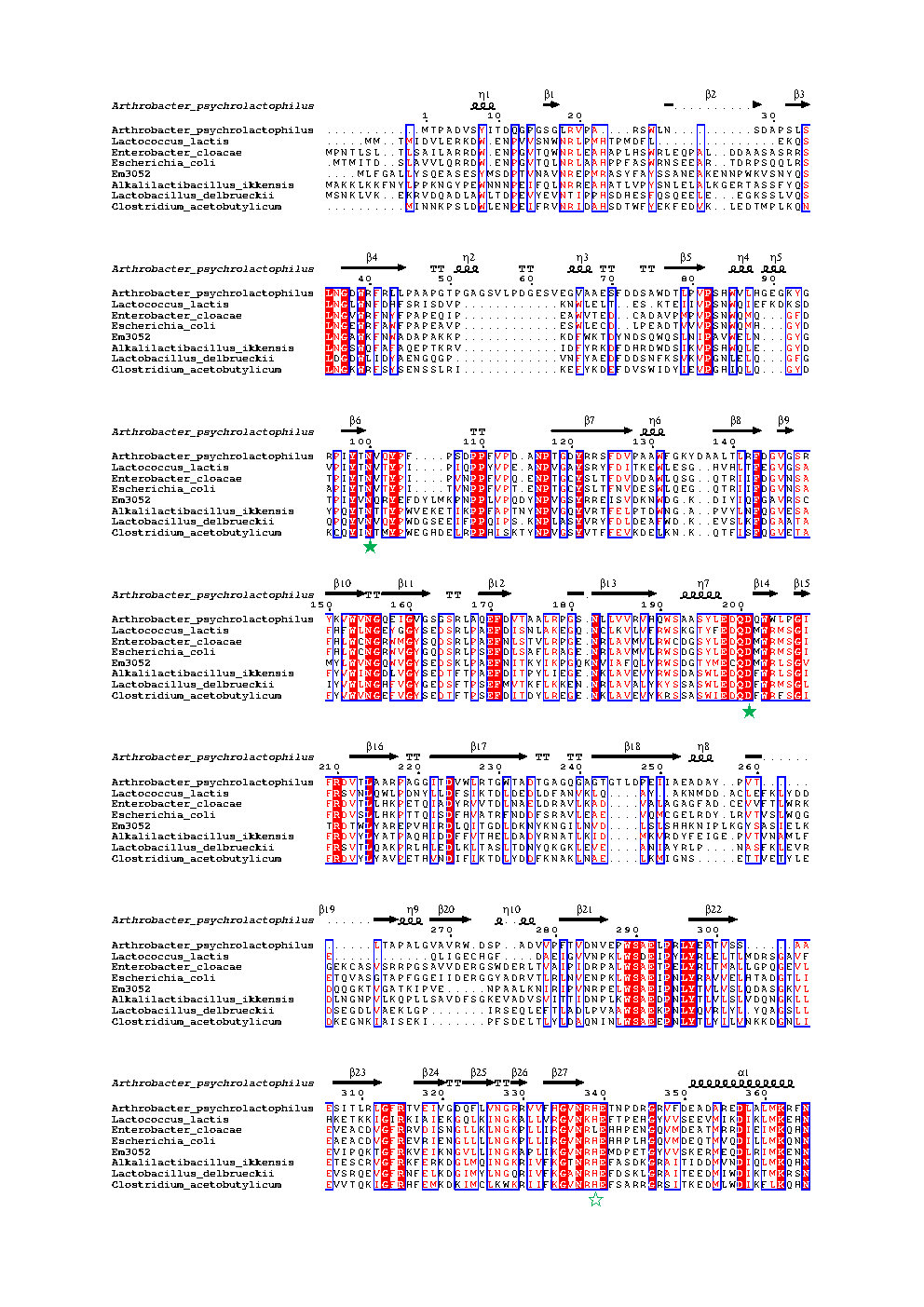

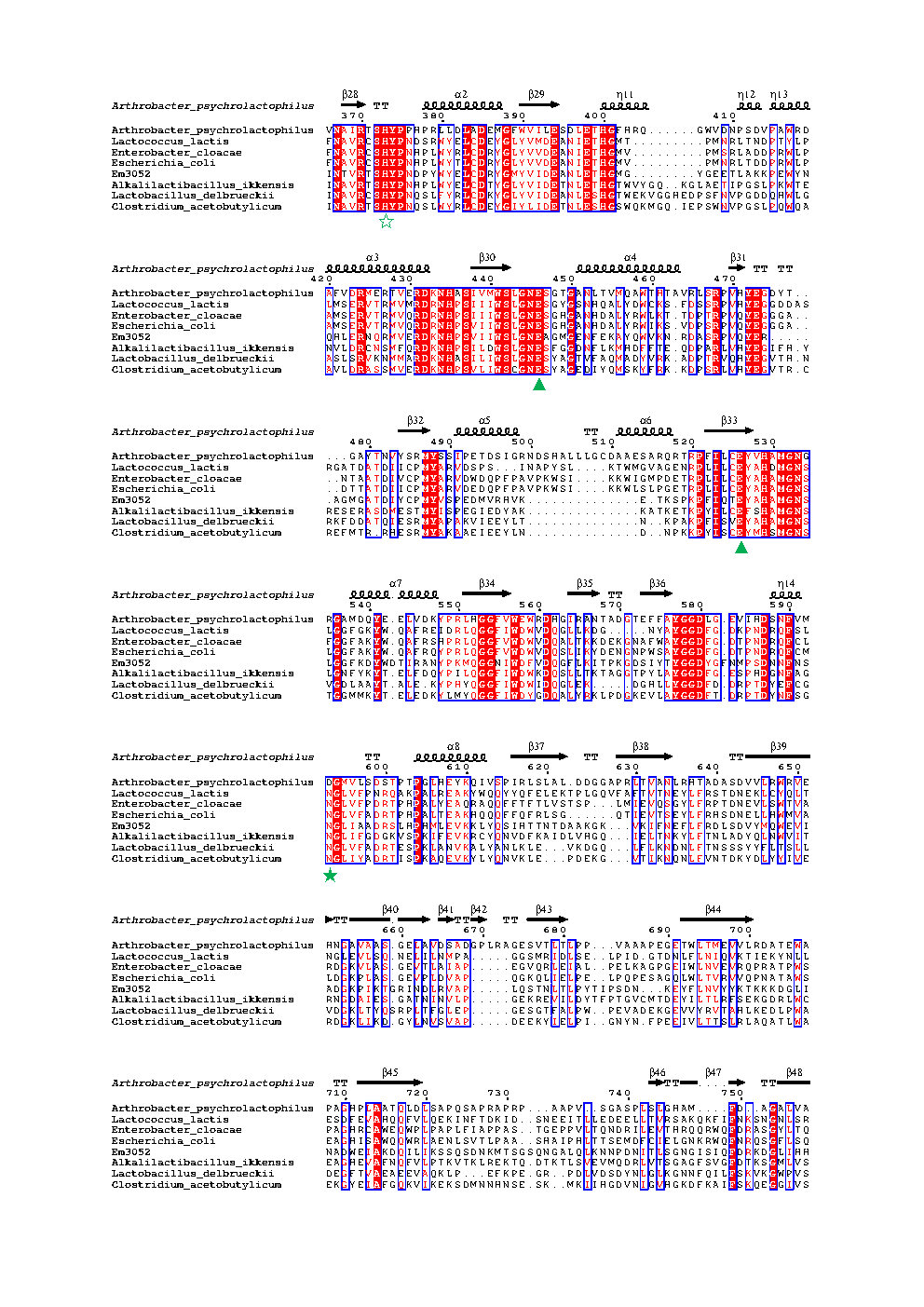

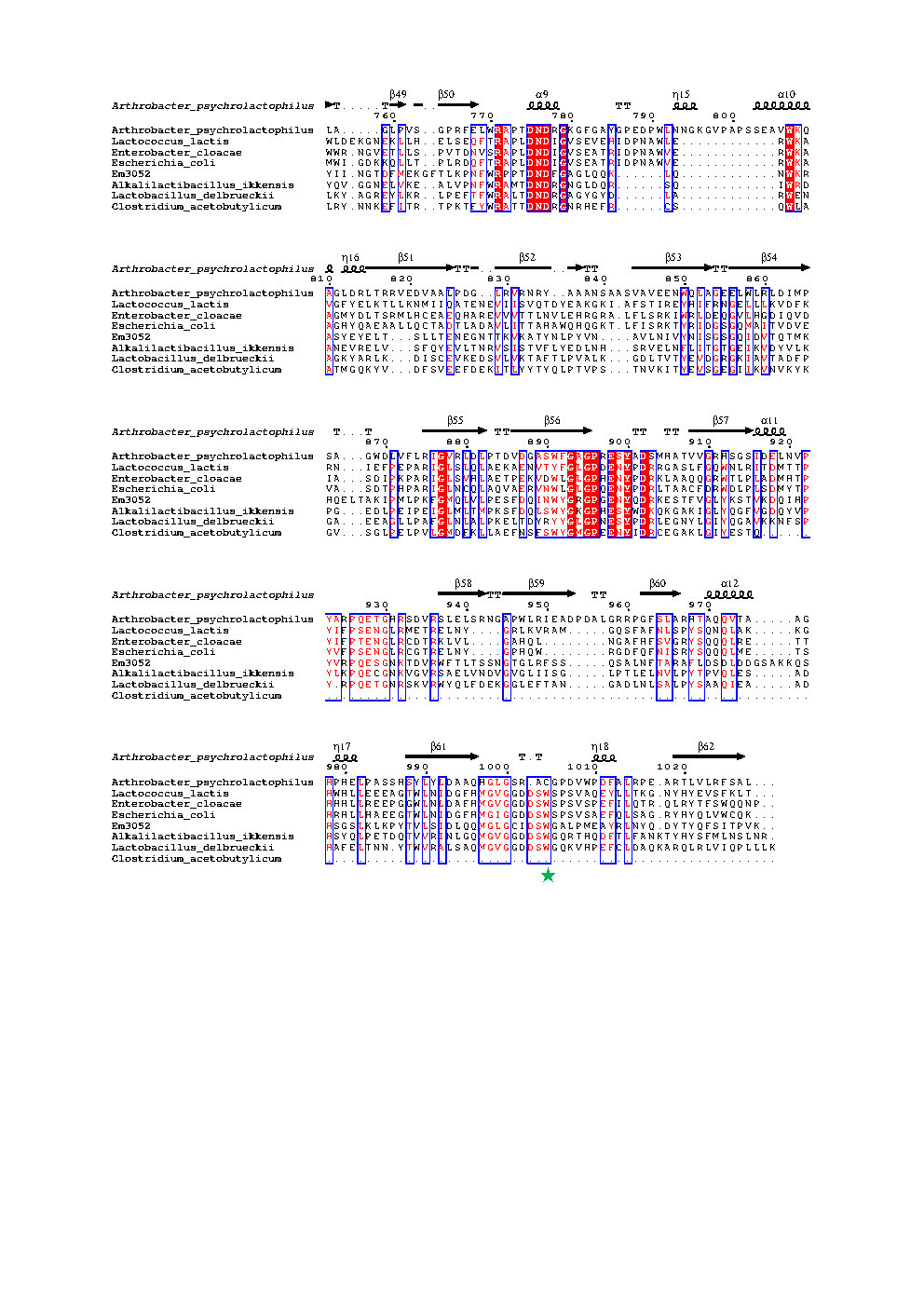

**Fig. S2 Sequence analysis of geGalaseI.** Structure-based sequence alignment of some GH2 family members. Through multiple sequence alignment, the potential binding sites (marked as green solid pentagram), stabilizers of the transition state (marked as green hollow pentagram) and potential catalytic sites (marked as green solid triangle) were predicted.
